# Evolution of mate harm resistance in females from *Drosophila melanogaster* populations selected for faster development and early reproduction

**DOI:** 10.1101/2022.12.25.521905

**Authors:** Tanya Verma, Susnato Das, Saunri Dhodi Lobo, Ashish Kumar Mishra, Soumi Bhattacharyya, Bodhisatta Nandy

**Author notes:** Corresponding author, email address, ORCID iD: 0000-0002-9588-0316. **Authors Contribution:** BN and TV conceptualized the study, designed the experiments, analysed and interpreted the results, and prepared the manuscript. SD helped with some parts of manuscript writing at earlier stage. TV, SD, SDL, AKM, SB executed the experiments, including data collection.

## Abstract

Interlocus sexual conflict is predicted to result in sexually antagonistic coevolution between male competitive traits, which are also female-detrimental, and mate harm resistance (MHR) in females. Little is known about connection life-history evolution and sexually antagonistic coevolution. Here, we investigated the evolution of MHR in a set of experimentally evolved populations, where mate-harming ability has been shown to have evolved in males as a correlated response to the selection for faster development and early reproduction. We measured mortality and fecundity of females of these populations and those of their matched controls, under different male exposure conditions. We observed that the evolved females were more susceptible to mate harm - suffering from significantly higher mortality under continuous exposure to control males within the twenty-day assay period. Though these evolved females are known to have shorter lifespan, such higher mortality was not observed under virgin and single-mating conditions. We used fecundity data to show that this higher mortality in evolved females is unlikely due to cost of egg production. Further analysis indicated that this decreased MHR is unlikely to be due purely to the smaller size of these females. Instead, it is more likely to be an indirect experimentally evolved response attributable to the changed breeding ecology, and/or male trait evolution. Our results underline the implications of changes in life history traits, including lifespan, to the evolution of MHR in females.

## Introduction

Evolutionary interests of sexes are often not aligned leading to evolutionary conflict over traits with sexually antagonistic fitness effects (Arnqvist & Rowe, 2005; Hosken et al., 2019; Parker, 1979). In one form of such conflict, commonly referred to as interlocus sexual conflict, expression of male-benefitting traits (for example, courtship and mating behavioural traits) reduces female fitness as an incidental side effect (Arnqvist & Rowe, 2005; Holland & Rice, 1999; Pitnick & García–González, 2002). The theory further predicts a counter evolution in female traits that reduce such mate harm – potentially resulting in sexually antagonistic coevolution (Arnqvist & Rowe, 2002; Dougherty et al., 2017; Friberg, 2005; Holland & Rice, 1998, 1999; Rankin et al., 2011; Snow et al., 2019; Wigby & Chapman, 2004). While interlocus conflict has been reported in a wide diversity of animals in the form of mate harm (Arnqvist, 1989; Chapman et al., 1995; Crudgington & Siva-Jothy, 2000; Fowler & Partridge, 1989; Partridge et al., 1987; Partridge & Fowler, 1990; Pitnick & García–González, 2002; Rice, 1996), direct observation of sexually antagonistic coevolution has been relatively difficult. Fruit fly *Drosophila. melanogaster* is an exceptional experimental system, where both interlocus conflict and sexually antagonistic coevolution are well studied. In this system, exposure to males is known reduce female fitness due to persistent courtship and mating attempt (von Schilcher & Dow, 1977), and also due to the side effects of the seminal fluid peptides received during a copulation (Chapman et al., 1995; Rice, 2000; Wolfner, 1997). These detrimental effects on females are expressed as increased mortality or lifetime reproductive output or both (Chapman et al., 1995; Holland & Rice, 1999; MacPherson et al., 2018; Nandy et al., 2013b, 2013a; Pitnick, 2001; Wigby & Chapman, 2004). There are now ample evidences, most notably from experimental evolution, showing the counter evolution of female resistance traits (Nandy et al., 2013c; Wigby & Chapman, 2004). In absence of the evolution of female resistance, hereafter referred to as mate-harm resistance (MHR), a population could suffer from reduced mean fitness potentially leading to extinction through the hitherto proposed tragedy-of-commons model (Rankin et al., 2007). Hence, understanding the causes and constraints relevant to the evolution of MHR is important.

MHR can include behavioural and physiological traits such as, avoidance of male encounter, finding refuges, production of proteins and peptides that respond to male seminal fluid protein (Arnqvist & Rowe, 2002; Chapman, 2018; Dougherty et al., 2017; Hopkins & Perry, 2022; Rice, 2000; Yun et al., 2017). These are expected to be physiologically costly for females. Though, to the best of our knowledge there hasn’t been any direct evidence or measure of such costs, there is some indirect evidence. For example, multiple experimental evolution results suggest reduction of MHR in females and the resulting increase in susceptibility of them to mate harm when populations were evolved under reduced sexual conflict (Holland & Rice, 1999; Wigby & Chapman, 2004).

Life history theories predict investment in costly traits such as, MHR should reduce when there is no fitness advantage of expressing such traits, especially when resources are constrained and/or there are stronger fitness components where resources are invested (Adler & Bonduriansky, 2014; Bonduriansky et al., 2008; Lemaître et al., 2020; Maklakov et al., 2007). These theories predict a trade-off between conflict related traits and somatic maintenance. Faster aging populations are thus expected to have greater investment in conflict related traits and vice versa (Promislow, 2003). Further, selection for life history that results in overall reduction in baseline resource availability, such as, selection for faster development, may also constrain the expression of conflict related traits (De Jong & Van Noordwijk, 1992; Van Noordwijk & De Jong, 1986).

It appears that relationship between life history and sexual selection/conflict is typical eco-evolutionary feedback (Bonduriansky, 2014; Rankin & Kokko, 2006), wherein ecological changes drive the evolution of reproductive and sexual traits through trade-offs and other phenotypic and genetic correlations culminating in changes in the breeding ecology. Since, intensity of sexual selection/conflict is a function of this can further change sexually selected traits. Such changes may further drive the evolutionary changes in life history traits. Recent experimental investigations examining the effect of evolution of faster development and early reproduction on the evolution of conflict related traits have upheld this idea. The relationship between life history and conflict related traits were found to be far more nuanced (Ghosh & Joshi, 2012; Mital et al., 2021, 2022; Verma et al., 2022).

In a previous report we showed evolutionary reduction in mate harming ability of *D. melanogaster* males is a set of populations subjected to the selection for faster development and early reproduction (Verma et al., 2022). Our results and those reported by Ghosh and Joshi (2012) and Mital et al. (2022) suggests that part of the changes in male traits can be attributable to the changes in key life history traits such as size. However, beyond the size effect, breeding ecology changes also play a clear role (Verma et al., 2022). It is, however, not clear if MHR in females respond to such changes in male traits. This is the question we address in the current manuscript.

Here we used four replicates of faster developing and early reproducing *D. melanogaster* populations (ACOs), and their controls (COs), to address this issue. We have previously reported the reduction of mate harming ability in ACO males. Therefore, we predicted that expression of MHR in ACO females to have no fitness advantage. Here we compare MHR in experimental (ACO) and control (CO) females to assess the evolution of MHR as a result of selection for faster development and early reproduction. We predict that if investment in sexually antagonistic traits is costly, experimental females should have reduced MHR. We measured MHR by assaying female mortality and fecundity under (a) virgin, (b) single mating and (c) continuous male exposure.

## Materials and methods

We used a set of experimentally evolved *D. melanogaster* populations. They consist five replicates of evolved populations, named ACOs, derived from five replicates of control population named COs. Detailed information on these populations can be found in the chapter 2. A total of eight populations consisting of four replicates of evolved ACO and their matched control CO populations were used for the present study. Hence, all assays described below were conducted with ACO_1_, ACO_2_, ACO_3_, ACO_4_ and their paired control CO populations. Each ACO population and their matched CO population have been treated together as one block in the present study. Hence, the assays were carried out in four distinct blocks where each block consisted of a replicate set of ACO and CO populations. All experimental flies were generated from a subset of the stock populations, after one generation of common garden rearing. All adult flies used for the experiments were collected as virgins. To obtain virgin flies, freshly eclosed flies were collected every 4-6 hours under light CO_2_ anaesthesia. Virgin flies were then held in single-sex vials at a density of 10 flies per vial until the assay. For a population, a total of 45 such vials of virgin females were collected. An adequate number of male vials were also collected from the corresponding CO population.

### Mate harm resistance assay setup

In *D. melanogaster*, MHR can be measured by comparing female mortality under limited and extended exposure to males (Jiang et al., 2011; Nandy et al., 2013c; Wigby & Chapman, 2004). Females with lower MHR are expected to show sharper increase in mortality under extended male exposure compared to those with higher MHR.

Assay vials were set up with 1-2 day old virgins. Each replicate population consisted of 45 vials, each vial having ten virgin females from a population, were randomly assigned to three assay conditions - virgin, single exposure, and continuous exposure such that each assay condition consisted of an initial count of 15 vials. The experimental vials were set up by introducing flies in fresh food vials. For the virgin assay condition, females were held without any male exposure for the entire assay duration. Single exposure and continuous exposure vials were set up by introducing 10 virgin control (i.e., CO) males along with the ten experimental females in a fresh food vial. We used control regime males (i.e., CO males) for this purpose to equalise the male background against which MHR of the evolved and control females was measured. For the single exposure vials, matings were manually observed and after a single round of mating, sexes were separated under CO_2_-anaesthesia to discard the males. Since under single exposure condition females received the mating exposure of males only once hence the single round of mating was conducted only once on assay set-up day.

After discarding the males the females were then returned back in the same vials. For the continuous exposure treatment, males and the females were kept together in the same vials till the end of the assay. To ensure similar handling of flies across all treatments, flies under virgin and continuous exposure treatments were also exposed to anaesthesia. Throughout the experiment, except sorting of sexes, all other fly handling was done without anaesthesia. All vials were maintained for twenty days and the flies in each vial were flipped to fresh food vials every alternate day. For all vials regardless of assay condition, mortality in females was recorded daily until day 20. Our previous observation suggests that the difference in effects of mate harm on female mortality can be detected in the first twenty days of adult life (Verma et al., 2022). In addition, this period represents early-to-mid-life in this system, most relevant to both control (CO) and experimental (ACO) population ecology. Further, the difference in age-dependent mortality rate between the two selection regimes has minimal impact on mortality difference within this duration (data not shown). Dead flies were aspirated out during vial-to-vial flips. In the continuous exposure assay condition, in case a female fly was found dead in a vial, along with the dead female, a male was also removed from the same vial to maintain a 1:1 sex ratio.

Female fecundity was recorded twice a week starting from the onset of the assay until day 20 (i.e., day 1, 3, 6, 9, 12, 15, 18, and 20). On each of these days, flies were flipped to a fresh food vial (hereafter referred to as a fecundity vial) and were left undisturbed for 24 hours.

Following this, the flies were transferred to a fresh food vial, while the fecundity vial was frozen immediately to prevent further development of the already deposited eggs. The number of eggs laid in a fecundity vial was counted under microscope. Fecundity count was carried out for single exposure and continuous exposure treatments. Per capita fecundity, calculated as total number of eggs in a vial divided by the number of females alive in that vial on that given day, from individual vials was taken as the unit of analysis. A few vials were removed from the assay for a variety of reasons, including accidental escape, a few females failing to mate, etc. The final sample size throughout the entire experiment was 13-15 vials per population.

### Data analysis

Female survivorship was analysed using Cox’s Proportional hazards model. Selection regime (levels: ACO and CO) and assay condition (levels: virgin, single exposure and continuous exposure) were modelled as fixed factor and block as random factor using R package `Coxme` (Therneau, 2012). Cox partial likelihood (log-likelihood) estimates across selection regimes were compared.

Per capita fecundity was analysed in two ways. Cumulative fecundity i.e., per capita fecundity pooled across all eight age classes was analysed to compare to total early-to-mid-life reproductive output of the females. In addition, age-specific per capita fecundity was analysed to compare the age-related pattern of reproduction. The latter was done only for the continuous exposure set to minimise model complication. Age-specific fecundity data were square root transformed before analysis. A linear mixed effect model was fitted to the transformed data. `lme4` package (Bates et al., 2015) and `lmerTest` (Kuznetsova et al., 2017) in R version 4.2.1 (R Core Team, 2022). In the cumulative fecundity model, selection regime (levels: ACO and CO), assay condition (levels: single exposure and continuous exposure) and their two-way interactions as fixed factors, block as a random factor. In the analysis of age-specific per capita fecundity, selection regime and age (levels: 1, 3, 6, 9, 12, 15, 18, 20) were the fixed factors, and block and all interaction terms involving block were modelled as random factors. All models are mentioned in the supplementary information.

Post-hoc pairwise comparisons using Tukey’s HSD method were performed with the package `Emmeans` (Lenth, 2016). The ANOVA table was obtained following Satterthwaite’s method using type III sum of squares.

## Results

Cox partial likelihood estimates suggested that the effects of selection regime, assay condition, and selection regime × assay condition interaction on female mortality were significant (Figure 1, Table 1). Pairwise comparisons indicated a significant difference in survivorship of ACO and CO females only under continuous exposure, with ACO females more than 9.5 times likely to succumb compared to CO females (estimated hazard ratio: 8.02, 95% CI: 3.786 to 17.798). ACO females are significantly smaller in size than CO females (see supplementary information for details). Therefore, to further investigate whether the higher mortality in ACOs compared to COs under CE condition was due to reduction in body size, we performed mortality analysis with and without thorax length (a proxy for body size) as a covariate. Details of the analysis can be found in supplementary information. The results of this analyses suggested that incorporating thorax length did not qualitatively change the interpretations of our results.

**Figure 1:**
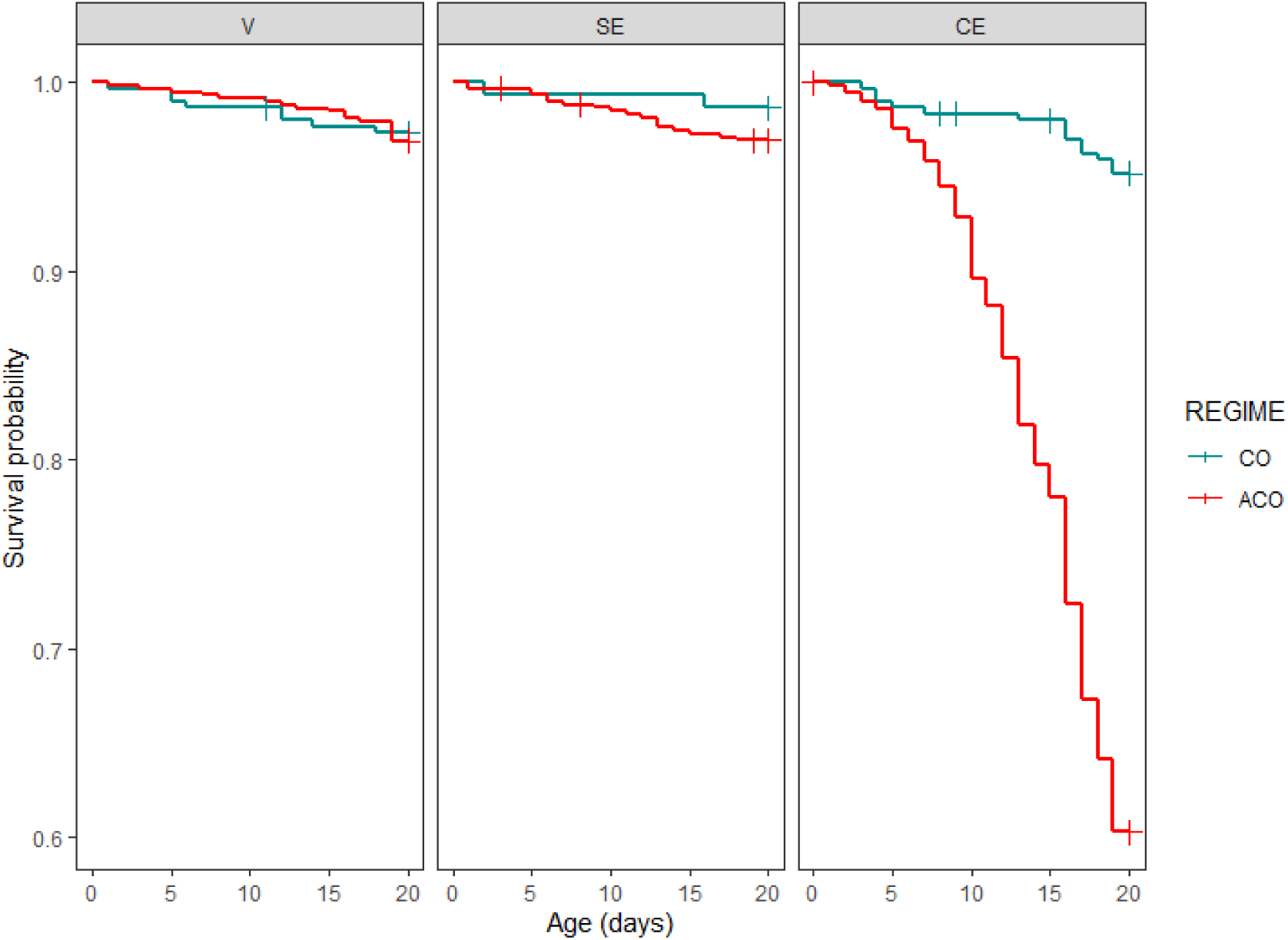
Survivorship curves obtained from Cox proportional hazard analysis on the mortality of ACO (red line) and CO (dark cyan line) regime females held under virgin (V), single exposure (SE) and continuous exposure (CE) condition for 20 days during the assay. The differences between survivorship ACO and CO females were found to be no significant under virgin and SE conditions. Under CE condition, ACO females showed significantly higher mortality rate.

**Table 1:** Output of mixed effect Cox proportional hazard model for analysis of female survivorship ACO and CO regime females held under virgin (V), single exposure (SE) and continuous exposure (CE) condition with ancestral CO males. Hazard ratios are relative to the default level for each factor which is set to 1. The default level for selection regime was ‘CO’, and the default level for assay condition was ‘Virgin’. Lower CI and Upper CI indicate lower and upper bounds of 95% confidence intervals. Level of significance was considered to be α = 0.05, and significant p-values are mentioned in bold font style

| **Fixed Coefficients** | **Hazard Ratios** | **Lower CI** | **Upper CI** | **z** | **p** |
| --- | --- | --- | --- | --- | --- |
| Selection Regime ACO | 1.156 | 0.589 | 2.266 | 0.24 | < **0.001** |
| Assay condition SE | 0.557 | 0.246 | 1.261 | -1.13 | < **0.001** |
| Assay condition CE | 1.927 | 1.051 | 3.536 | 1.36 | < **0.001** |
| Selection Regime ACO: Assay condition SE | 1.776 | 0.624 | 5.053 | 0.98 | < **0.001** |
| Selection Regime ACO: Assay condition CE | 8.209 | 3.786 | 17.798 | 4.27 | < **0.001** |
| **Random effects** | **Variance** |  |  |  |  |
| Block | 0.0794 |  |  |  |  |

The effects of selection regime and assay condition on cumulative fecundity were significant (Table 2). While females under continuous exposure had significantly higher fecundity regardless of the selection regime, cumulative fecundity of ACO females was 27% less than that of the control CO females (Figure 2a). Age-specific fecundity analysis indicated significant effects of selection regime, and age (Figure 2b). However, we found a two-way and a three-way interaction term involving random block to be significant (see SI, Table S3). Hence, we analysed each block separately (see supplementary information, Table S2).

**Figure 2:**
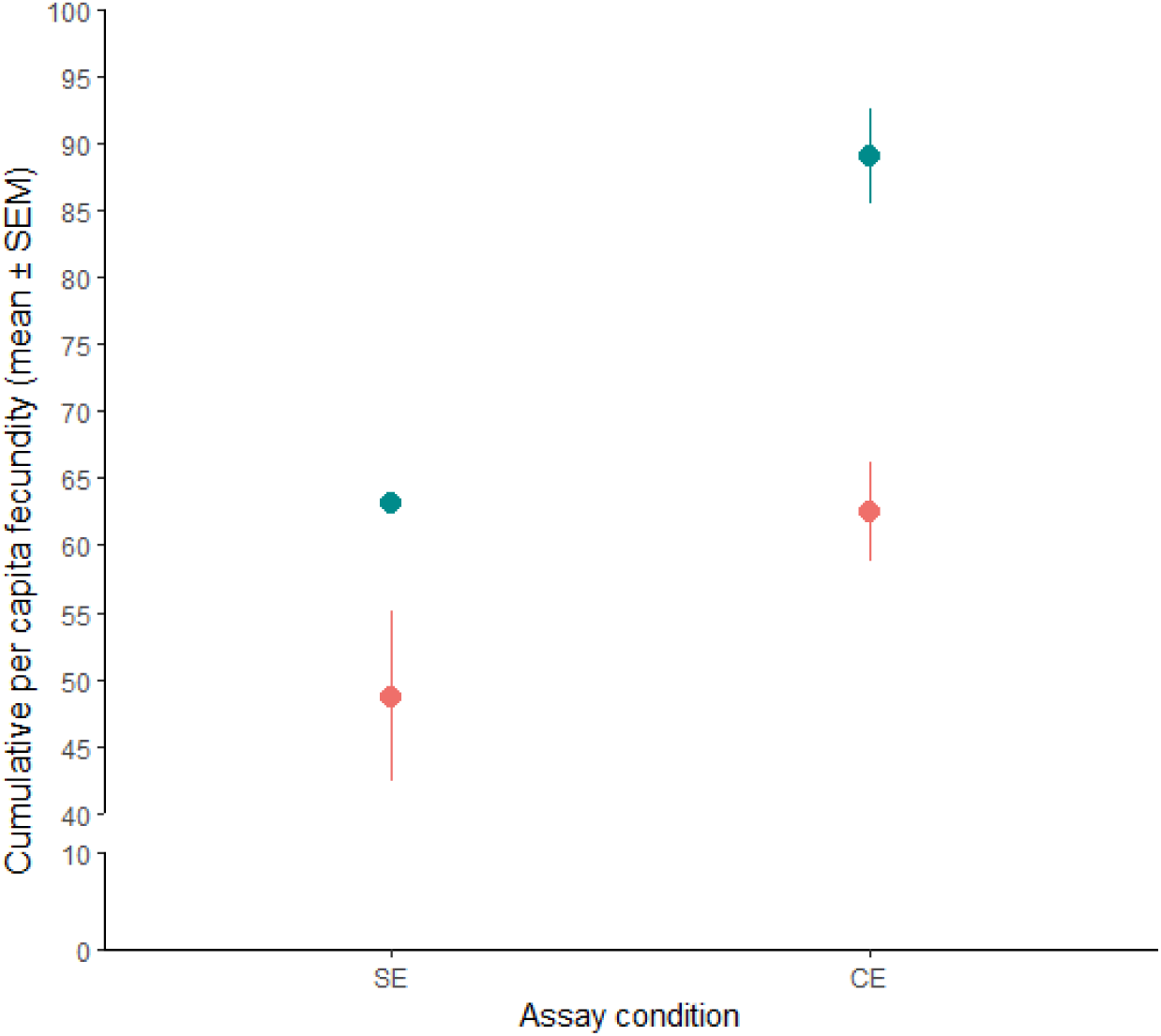
Results from the cumulative fecundity per capita across ACO and CO selection regime females held with control (CO) males. Filled circles and error bars represent means, and standard error respectively. Standard errors are calculated using block means (i.e., population means). Effects of selection regime, and assay condition on cumulative per capita fecundity were found to be significant.

**Figure 3:**
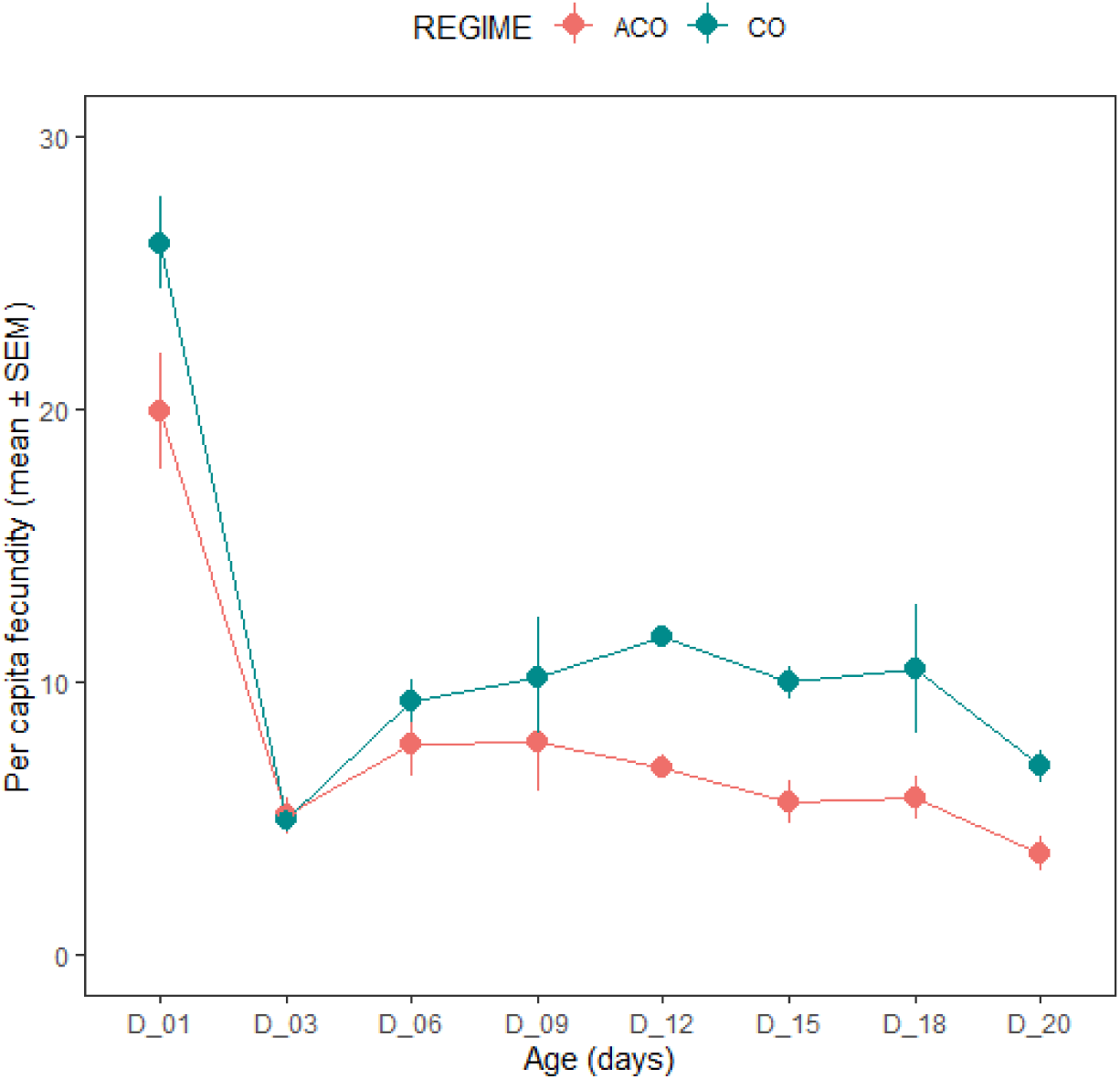
Results from the age-specific per capita fecundity across ACO and CO selection regime females held with control (CO) males. Age specific fecundity was analysed only for continuous exposure assay condition to minimise model complication. Filled circles and error bars represent means, and standard error respectively. Standard errors are calculated using block means (i.e., population means). Effects of selection regime and age were found to be significant on age specific per capita fecundity.

**Table 2:** Summary of the results of linear mixed model (LMM) analysis of cumulative fecundity, day 1 per capita fecundity, age-specific per capita fecundity and body size using `lmerTest` function in R. Selection regime and assay condition in cumulative and day 1 per capita fecundity and regime and Selection regime in body size were modelled as fixed factors and block as a random factor. All tests were done considering α = 0.05 and significant p-values are mentioned in bold font style.

| Trait | Effect | SS | DF | MS | Den<br>DF | F | p |
| --- | --- | --- | --- | --- | --- | --- | --- |
| **Cumulative**<br>**fecundity** | Selection Regime (SR) | 4054.8 | 1 | 4054.8 | 3.008 | 25.826 | 0.015 |
|  | Assay condition | 5476.4 | 1 | 5476.4 | 2.987 | 34.881 | 0.010 |
|  | Selection Regime ×<br>Assay condition | 583.0 | 1 | 583.0 | 3.016 | 3.713 | 0.149 |
| **Day 1 per capita**<br>**fecundity** | Selection Regime (SR) | 301.22 | 1 | 301.22 | 5.649 | 11.673 | **0.016** |
|  | Assay condition | 296.34 | 1 | 296.34 | 4.470 | 11.484 | **0.023** |
|  | Selection Regime ×<br>Assay condition | 129.31 | 1 | 129.31 | 2.077 | 5.011 | 0.150 |
| **Age-specific per**<br>**capita fecundity** | Selection regime | 14.20 | 1 | 14.21 | 40.027 | 42.210 | **<0.001** |
|  | Age | 48.57 | 7 | 6.94 | 21.019 | 20.626 | **<0.001** |
|  | Selection regime × Age | 5.19 | 7 | 0.74 | 23.470 | 2.205 | 0.071 |
| **Body size** | Selection Regime | 1.11 | 1 | 1.11 | 235.000 | 1446.600 | **<0.001** |

Though across blocks the age-specific pattern seemed to vary, CO females generally showed higher per-capita fecundity in most age points (see SI, Figure S3). Fecundity on day1 was of particular interest as ACO maintenance regime selects for fecundity at this age. Hence, we analysed day1 fecundity separately, using a linear mixed model similar to that used to analyse cumulative fecundity. The results indicated significant effects of selection regime and assay condition, with CO females showing higher fecundity under both SE and CE conditions (Table 6.2, Figure S4).

## Discussion

Our results suggest that selection for faster development and early reproduction has led to the evolution of sexually antagonistic traits. The evolved ACO males were previously shown to be significantly less harming to their mates (Verma et al., 2022). According to the results of our MHR assay reported here, ACO females appeared to be significantly more susceptible to continued male interaction. When held with control males, ACO females showed close to ten times higher mortality rate compared to that of the control (CO) females in the same condition. Further, ACO females were found to be consistently less fecund, regardless of the length of male exposure, and age. Hence, higher mortality of ACO females mentioned above was more likely to be due to an increased susceptibility to male induced harm, instead of an increased cost of reproduction *per se*.

Experimental evolution resulted in reduced adult longevity in the ACOs, and females are known to have >30% reduced mean lifespan (Burke et al., 2010). Hence, ACO females are expected to have a higher mortality. However, no difference in survivorship of females were observed when they were kept under absence of male exposure i.e., virgin condition or even under single mating exposure for the entire twenty-day assay period. This shows that there is no intrinsic difference in the mortality rate of females of two selection regime, at least for the assay period. Therefore, higher mortality under continued presence of males represents mortality due to male induced harm and thus not the intrinsic mortality differences between selection regime females. Reduction in MHR as a coevolutionary response to the reduction in mate harm from males should depend on the cost of maintaining and/or expressing MHR traits. Indeed, under reduced level of sexual conflict experimental evolution have repeatedly led to the reduction in MHR (Nandy et al., 2013c; Wigby & Chapman, 2004).

The harm induced by males to female is often measured in terms of reduced survival rate. In *Drosophila*, males are known to reduce female survival through (a) detrimental effect of seminal fluid proteins (Sfp) transferred during copulation (Chapman et al., 1995; Wigby et al., 2020) and (b) by persistently courtship to the females (Fowler & Partridge, 1989). Several experimental evolution studies have previously shown the evolution of female resistance to male induced harm (Crudgington et al., 2005; Holland & Rice, 1999; Maklakov et al., 2007; Martin & Hosken, 2003; Michalczyk et al., 2011; Nandy et al., 2013c, Wigby & Chapman, 2004). Reduction in population level sexual conflict, either through enforced monogamy or through altered operational sex ratio emerged as the fundamental selective condition needed for such female evolution (for example, Martin & Hosken, 2003, Nandy et al., 2013c). Since ACO males are already shown to be less harming than ancestral CO males (Verma et al., 2022), it is perhaps reasonable to suggest that ACO females are usually subjected to much less sexually antagonistic male interactions. Hence, the female resistance traits, in absence of any selective advantage, are free to evolve due to their costs.

If MHR is costly to express, it is expected to be constrained by the resource availability (Adler & Bonduriansky, 2014). Females in resource deprived condition should therefore be limited in terms of their ability to resist mate harm. Such condition dependence of MHR has been recently demonstrated (Iglesias-Carrasco et al., 2018; Rostant et al., 2020). In addition, for reproducing females, the cost of producing progeny can further constrain resources available for other physiological processes - potentially making them vulnerable to stresses including mate harm. The evolved ACO females in our study are small in size (Table 2 and see Supplementary information), and can thus be expected to be resource limited (Chippindale et al., 1993). However, they have a lower reproductive rate - hence, lower absolute investment in reproduction. Though it is difficult to assess the relative reproductive investment, as evident from our data from the single mating treatment, there appears to be a baseline reduction in reproductive rate of ACO females. However, evidently this baseline difference in reproduction did not result in mortality rate difference, which is only evident under extended male exposure. In addition, there was no evidence that this difference in reproductive investment between the evolved ACO and control CO females was higher under continuous male presence. Hence, it is very unlikely that observed differences in susceptibility is a mere reflection of the difference in available resources after accounting for the resources needed for reproduction *per se*. The size difference, however, could still be a fundamental reason for reduction of MHR of the ACO females. Notwithstanding the potential effect of body size on our MHR interpretation, re-analysis of the mortality results with female thorax length as a covariate did not qualitatively change final outcome of the analysis. Further, our conclusions are also in line with those of Mital et al. (2021) who used phenocopied females to demonstrate the size independent reduction in MHR. The literature is also fairly ambivalent about the dependence of MHR on female size. Hence, in conclusion, it is very unlikely that the reduction in MHR of the ACO females can be completely attributable to reduced size of these females, however, it cannot be completely ruled out either.

Several experimental evolution studies have shown the evolution of MHR (Crudgington et al., 2005; Holland & Rice, 1999; Hollis et al., 2019, p. 2020; Maklakov et al., 2007; Martin & Hosken, 2003; Michalczyk et al., 2011; Mital et al., 2021; Nandy et al., 2013c; Rostant et al., 2020; Wigby & Chapman, 2004). Of these, only two have directly connected evolution of conflict related traits to life history traits such as, condition, adult lifespan, development time, and size (Mital et al., 2022; Rostant et al., 2020). Though evolution of MHR is important for a population’s survival (Rankin et al., 2011), continuation of sexual selection (Snow et al., 2019), and maintenance of genetic variation (Härdling & Karlsson, 2009), it cannot evolve in the vacuum of sexually antagonistic traits only. Our results are an important addition to the growing list of evidences suggesting that sexual conflict is subjected to a typical eco-evolutionary feedback process. A key prediction of this is, the ecological variations such as population dynamics, competition, economics of mating interactions can drive the evolutionary changes (i.e. “eco-evo” dynamics) and evolutionary changes can in turn influence the ecological processes such as population dynamics, productivity, investment etc. (“evo-eco” dynamics: (Svensson, 2018)). Within-population studies showed that ecological factors such as availability of food, predation pressure, operational sex ratio can affect mating economics and interactions thereby affecting the degree of sexual selection and sexual conflict (Ortigosa & Rowe, 2002; Perry & Rowe, 2018). For example, Ortigosa and Rowe (2002) showed that in water strider (*Gerris buenoi*) females under low availability of food, increases behavioural resistance to mating because mating interferes with female foraging, thereby strengthening sexual selection to favour male investment in persistence traits. Additionally, across population studies have also supported the idea that ecological variations affecting male-female encounter can be an important determinant of investment in sexually antagonistic traits. For example, complexity of the mating environment providing refuges for mate avoidance (Byrne et al., 2008; Yun et al., 2017, 2021), temperature variation that alters various activities in males (García-Roa et al., 2019), and community structure that alters male-female encounter rate (Clutton-Brock et al., 1999; Gomez-Llano et al., 2018) have been found to alter the level of mate harm in a population. Therefore, breeding ecology can set the stage of sexual conflict and drive antagonistic coevolution between sexes, and moreover, life history can affect such evolution by (a) setting physiological and genetic constraints, and (b) constraining breeding ecology. Hence, selection for life history traits such as, lifespan, reproductive schedule etc. should be important drivers of sexually antagonistic coevolution as such selection can impact breeding ecology and offset the fitness premium on sexually antagonistic traits.

In conclusion, we found that the reproductive evolution in the ACO females. The results suggested that as a correlated response to the selection for faster development and early reproduction, female fecundity and resistance to mate harm had evolved. Much of the changes in resistance trait can be attributed to the incidental changes in the breeding ecology in addition to the potential effect of resource limitation.

## Acknowledgements

The study was financially supported by a research grant from Department of Science and Technology, Govt. of India (INSPIRE Faculty award, Grant no. DST/INSPIRE/04/2013/000520). We thank Subhasish Halder and Purbasha Dasgupta for help in the experiments and Rabisankar Pal for help in data analysis and plotting. We thank Sadanjeet Kumar Kar for help in experimental observations. TV thanks Indian Institute of Science Education and Research, Berhampur for financial support in the form of Junior and Senior Research Fellowship. SD thanks Scholarship for Higher Education, Govt. of India for financial support in form of INSPIRE-SHE fellowship. SDL thanks IASc, INSA and NASI, Government of India for financial support in the form of IASc-INSA-NASI Summer Research fellowship. AKM thanks Department of Atomic Energy, Govt. of India for financial support in the form of DISHA scholarship. We do not have any conflict of interest to declare.

## Notes

### Competing Interest Statement

The authors have declared no competing interest.

### Summary of Updates

The revision includes no new data, but some re-analyses, and new analysis. Much of the discussionn and abstract have been revised. The overall conclusion and the story have not changed in the revised version.

## References

Adler, M. I., & Bonduriansky, R. (2014). Sexual conflict, life span, and aging. Cold Spring Harbor Perspectives in Biology, 6(8), a017566.

Arnqvist, G. (1989). Sexual selection in a water strider: The function, mechanism of selection and heritability of a male grasping apparatus. Oikos, 344–350.

Arnqvist, G., & Rowe, L. (2002). Antagonistic coevolution between the sexes in a group of insects. Nature, 415(6873), 787–789. https://doi.org/10.1038/415787a

Arnqvist, G., & Rowe, L. (2005). Sexual conflict (Vol. 31). Princeton university press.

Bates, D., Kliegl, R., Vasishth, S., & Baayen, H. (2015). Parsimonious mixed models. ArXiv Preprint ArXiv:1506.04967.

Bonduriansky, R. (2014). The ecology of sexual conflict: Background mortality can modulate the effects of male manipulation on female fitness. Evolution, 68(2), 595–604.

Bonduriansky, R., Maklakov, A., Zajitschek, F., & Brooks, R. (2008). Sexual selection, sexual conflict and the evolution of ageing and life span. Functional Ecology, 443–453.

Burke, M. K., Dunham, J. P., Shahrestani, P., Thornton, K. R., Rose, M. R., & Long, A. D. (2010). Genome-wide analysis of a long-term evolution experiment with *Drosophila*. Nature, 467(7315), 587–590.

Byrne, P. G., Rice, G. R., & Rice, W. R. (2008). Effect of a refuge from persistent male courtship in the Drosophila laboratory environment. American Zoologist, 48(2), e1–e1.

Chapman, T. (2018). Sexual conflict: Mechanisms and emerging themes in resistance biology. The American Naturalist, 192(2), 217–229.

Chippindale, A. K., Leroi, A. M., Kim, S. B., & Rose, M. R. (1993). Phenotypic plasticity and selection in *Drosophila* life-history evolution. I. Nutrition and the cost of reproduction. Journal of Evolutionary Biology, 6(2), 171–193.

Chapman, T., Liddle, L. F., Kalb, J. M., Wolfner, M. F., & Partridge, L. (1995). Cost of mating in Drosophila melanogaster females is mediated by male accessory gland products. Nature, 373(6511), 241–244.

Clutton-Brock, T. H., Maccoll, A., Chadwick, P., Gaynor, D., Kansky, R., & Skinner, J. D. (1999). Reproduction and survival of suricates (Suricata suricatta) in the southern Kalahari. African Journal of Ecology, 37(1), 69–80.

Crudgington, H. S., Beckerman, A. P., Brüstle, L., Green, K., & Snook, R. R. (2005). Experimental removal and elevation of sexual selection: Does sexual selection generate manipulative males and resistant females? The American Naturalist, 165(S5), S72–S87.

Crudgington, H. S., & Siva-Jothy, M. T. (2000). Genital damage, kicking and early death. Nature, 407(6806), 855–856.

De Jong, G., & Van Noordwijk, A. J. (1992). Acquisition and allocation of resources: Genetic (co) variances, selection, and life histories. The American Naturalist, 139(4), 749–770.

Dougherty, L. R., van Lieshout, E., McNamara, K. B., Moschilla, J. A., Arnqvist, G., & Simmons, L. W. (2017). Sexual conflict and correlated evolution between male persistence and female resistance traits in the seed beetle Callosobruchus maculatus. Proceedings of the Royal Society B: Biological Sciences, 284(1855), 20170132.

Fowler, K., & Partridge, L. (1989). A cost of mating in female fruitflies. Nature, 338(6218), 760–761.

Friberg, U. (2005). Genetic variation in male and female reproductive characters associated with sexual conflict in *Drosophila melanogaster*. Behavior Genetics, 35(4), 455–462.

García-Roa, R., Chirinos, V., & Carazo, P. (2019). The ecology of sexual conflict: Temperature variation in the social environment can drastically modulate male harm to females. Functional Ecology, 33(4), 681–692.

Ghosh, S., & Joshi, A. (2012). Evolution of reproductive isolation as a by-product of divergent life-history evolution in laboratory populations of D rosophila melanogaster. Ecology and Evolution, 2(12), 3214–3226.

Gomez-Llano, M. A., Bensch, H. M., & Svensson, E. I. (2018). Sexual conflict and ecology: Species composition and male density interact to reduce male mating harassment and increase female survival. Evolution, 72(4), 906–915.

Gromko, M. H., & Pyle, D. W. (1978). Sperm competition, male fitness, and repeated mating by female Drosophila melanogaster. Evolution, 588–593.

Härdling, R., & Karlsson, K. (2009). The dynamics of sexually antagonistic coevolution and the complex influences of mating system and genetic correlation. Journal of Theoretical Biology, 260(2), 276–282.

Holland, B., & Rice, W. R. (1998). Perspective: Chase-away sexual selection: antagonistic seduction versus resistance. Evolution, 52(1), 1–7.

Holland, B., & Rice, W. R. (1999). Experimental removal of sexual selection reverses intersexual antagonistic coevolution and removes a reproductive load. Proceedings of the National Academy of Sciences, 96(9), 5083–5088.

Hollis, B., Koppik, M., Wensing, K. U., Ruhmann, H., Genzoni, E., Erkosar, B., Kawecki, T. J., Fricke, C., & Keller, L. (2019). Sexual conflict drives male manipulation of female postmating responses in Drosophila melanogaster. Proceedings of the National Academy of Sciences, 116(17), 8437–8444.

Hopkins, B. R., & Perry, J. C. (2022). The evolution of sex peptide: Sexual conflict, cooperation, and coevolution. Biological Reviews, 97(4), 1426–1448.

Hosken, D. J., Archer, C. R., & Mank, J. E. (2019). Sexual conflict. Current Biology, 29(11), R451–R455.

Iglesias-Carrasco, M., Jennions, M. D., Zajitschek, S. R., & Head, M. L. (2018). Are females in good condition better able to cope with costly males? Behavioral Ecology, 29(4), 876–884. https://doi.org/10.1093/beheco/ary059. https://doi.org/10.1093/beheco/ary059

Jiang, P.-P., Bedhomme, S., Prasad, N. G., & Chippindale, A. (2011). Sperm competition and mate harm unresponsive to male-limited selection in Drosophila: An evolving genetic architecture under domestication. Evolution: International Journal of Organic Evolution, 65(9), 2448–2460.

Kaufman, B. P., & Demerec, M. (1942). Utilization of sperm by the female Drosophila melanogaster. The American Naturalist, 76(766), 445–469.

Kuijper, B., Stewart, A. D., & Rice, W. R. (2006). The cost of mating rises nonlinearly with copulation frequency in a laboratory population of *Drosophila melanogaster*. Journal of Evolutionary Biology, 19(6), 1795–1802.

Kuznetsova, A., Brockhoff, P. B., & Christensen, R. H. (2017). lmerTest package: Tests in linear mixed effects models. Journal of Statistical Software, 82, 1–26.

Lemaître, J.-F., Ronget, V., & Gaillard, J.-M. (2020). Female reproductive senescence across mammals: A high diversity of patterns modulated by life history and mating traits. Mechanisms of Ageing and Development, 192, 111377.

Lenth, R. V. (2016). Least-squares means: The R package lsmeans. Journal of Statistical Software, 69, 1–33.

MacPherson, A., Yun, L., Barrera, T. S., Agrawal, A. F., & Rundle, H. D. (2018). The effects of male harm vary with female quality and environmental complexity in Drosophila melanogaster. Biology Letters, 14(8), 20180443.

Maklakov, A. A., Fricke, C., & Arnqvist, G. (2007). Sexual selection affects lifespan and aging in the seed beetle. Aging Cell, 6(6), 739–744.

Martin, O. Y., & Hosken, D. J. (2003). Costs and benefits of evolving under experimentally enforced polyandry or monogamy. Evolution, 57(12), 2765–2772.

Michalczyk, Lukasz, Millard, A. L., Martin, O. Y., Lumley, A. J., Emerson, B. C., & Gage, M. J. (2011). Experimental evolution exposes female and male responses to sexual selection and conflict in Tribolium castaneum. Evolution: International Journal of Organic Evolution, 65(3), 713–724.

Mital, A., Sarangi, M., Dey, S., & Joshi, A. (2021). Evolution of lower levels of inter-locus sexual conflict in D. melanogaster populations under strong selection for rapid development. BioRxiv.

Mital, A., Sarangi, M., Nandy, B., Pandey, N., & Joshi, A. (2022). Shorter effective lifespan in laboratory populations of D. melanogaster might reduce sexual selection. BioRxiv, 2021–04.

Nandy, B., Chakraborty, P., Gupta, V., Ali, S. Z., & Prasad, N. G. (2013a). Sperm competitive ability evolves in response to experimental alteration of operational sex ratio. Evolution, 67(7), 2133–2141.

Nandy, B., Gupta, V., Sen, S., Udaykumar, N., Samant, M. A., Ali, S. Z., & Prasad, N. G. (2013b). Evolution of mate-harm, longevity and behaviour in male fruit flies subjected to different levels of interlocus conflict. BMC Evolutionary Biology, 13(1), 1–16.

Nandy, B., Gupta, V., Udaykumar, N., Samant, M. A., Sen, S., & Prasad, N. G. (2013c). Experimental evolution of female traits under different levels of intersexual conflict in *Drosophila melanogaster*. Evolution, 68(2), 412–425. https://doi.org/10.1111/evo.12271

Ortigosa, A., & Rowe, L. (2002). The effect of hunger on mating behaviour and sexual selection for male body size in Gerris buenoi. Animal Behaviour, 64(3), 369–375.

Parker, G. A. (1979). Sexual selection and sexual conflict. Sexual Selection and Reproductive Competition in Insects, 123, 166.

Partridge, L., & Fowler, K. (1990). Non-mating costs of exposure to males in female *Drosophila melanogaster*. Journal of Insect Physiology, 36(6), 419–425.

Partridge, L., Green, A., & Fowler, K. (1987). Effects of egg-production and of exposure to males on female survival in Drosophila melanogaster. Journal of Insect Physiology, 33(10), 745–749.

Perry, J. C., & Rowe, L. (2018). Sexual conflict in its ecological setting. Philosophical Transactions of the Royal Society B: Biological Sciences, 373(1757), 20170418.

Pitnick. (2001). Evolution of female remating behaviour following experimental removal of sexual selection. Proceedings of the Royal Society of London. Series B: Biological Sciences, 268(1467), 557–563.

Pitnick, S., & García–González, F. (2002). Harm to females increases with male body size in Drosophila melanogaster. Proceedings of the Royal Society of London. Series B: Biological Sciences, 269(1502), 1821–1828.

Promislow, D. (2003). Mate choice, sexual conflict, and evolution of senescence. Behavior Genetics, 33(2), 191–201.

Rankin, D. J., Bargum, K., & Kokko, H. (2007). The tragedy of the commons in evolutionary biology. Trends in Ecology & Evolution, 22(12), 643–651.

Rankin, D. J., Dieckmann, U., & Kokko, H. (2011). Sexual conflict and the tragedy of the commons. The American Naturalist, 177(6), 780–791.

Rankin, D. J., & Kokko, H. (2006). Sex, death and tragedy. Trends in Ecology & Evolution, 21(5), 225–226.

Rice, W. R. (1996). Sexually antagonistic male adaptation triggered by experimental arrest of female evolution. Nature, 381, 232–234.

Rice, W. R. (2000). Dangerous liaisons. Proceedings of the National Academy of Sciences, 97(24), 12953–12955.

Rostant, W. G., Fowler, E. K., & Chapman, T. (2020). Sexual Conflict Theory: Concepts and Empirical Tests. In The Sage Handbook of Evolutionary Psychology: Foundations of Evolutionary Psychology (pp. 241–259). SAGE Publications.

Snow, S. S., Alonzo, S. H., Servedio, M. R., & Prum, R. O. (2019). Female resistance to sexual coercion can evolve to preserve the indirect benefits of mate choice. Journal of Evolutionary Biology, 32(6), 545–558.

Stearns, S. C. (1989). Trade-offs in life-history evolution. Functional Ecology, 3(3), 259–268.

Svensson. (2018). On reciprocal causation in the evolutionary process. Evolutionary Biology, 45(1), 1–14.

Therneau, T. (2012). Coxme: Mixed effects Cox models. R package. Version.

Van Noordwijk, A. J., & De Jong, G. (1986). Acquisition and allocation of resources: Their influence on variation in life history tactics. The American Naturalist, 128(1), 137–142.

Verma, T., Mohapatra, A., Senapati, H. K., Muni, R. K., Dasgupta, P., & Nandy, B. (2022). Evolution of reduced mate harming tendency of males in *Drosophila melanogaster* populations selected for faster life history. Behavioral Ecology and Sociobiology, 76(6), 1– 15.

von Schilcher, F., & Dow, M. (1977). Courtship behaviour in Drosophila: Sexual isolation or sexual selection? Zeitschrift Für Tierpsychologie, 43(3), 304–310.

Wigby, S., Brown, N. C., Allen, S. E., Misra, S., Sitnik, J. L., Sepil, I., Clark, A. G., & Wolfner, M. F. (2020). The Drosophila seminal proteome and its role in postcopulatory sexual selection. Philosophical Transactions of the Royal Society B, 375(1813), 20200072.

Wigby, S., & Chapman, T. (2004). Female resistance to male harm evolves in response to manipulation of sexual conflict. Evolution, 58(5), 1028–1037.

Wolfner, M. F. (1997). Tokens of love: Functions and regulation of Drosophila male accessory gland products. Insect Biochemistry and Molecular Biology, 27(3), 179–192.

Yun, L., Agrawal, A. F., & Rundle, H. D. (2021). On male harm: How it is measured and how it evolves in different environments. The American Naturalist, 198(2), 219–231.

Yun, L., Chen, P. J., Singh, A., Agrawal, A. F., & Rundle, H. D. (2017). The physical environment mediates male harm and its effect on selection in females. Proceedings of the Royal Society B: Biological Sciences, 284(1858), 20170424.

