## Supplementary information for "Evolution of mate harm resistance in females from *Drosophila melanogaster* populations selected for faster development and early reproduction"

**S1: Measurement of thorax length as a proxy for body size**

A separate set of 30 ACO females and 30 CO females was collected on eclosion for each population and frozen at -20 ⁰C for body size measurement. Thorax length was measured as a proxy for body size. Images of all the flies were taken later. These images were then used to measure the length of the thorax of each fly on the ImageJ software. The length was measured from the midpoint of the anterior margin of the thorax to the distal midpoint of the scutellum.

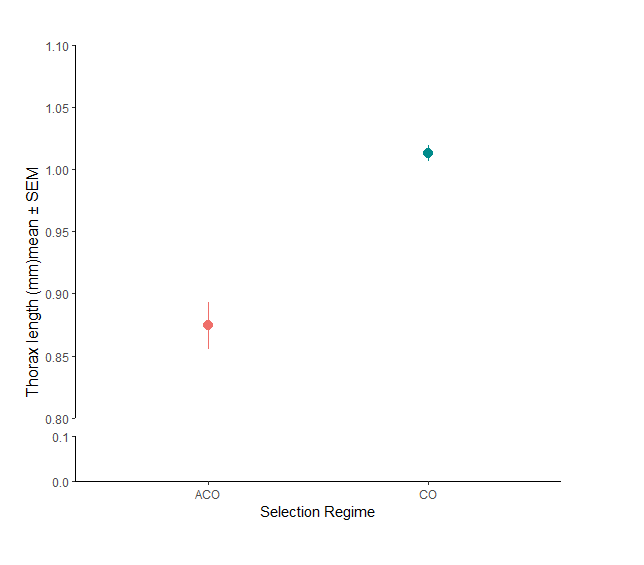

**Figure S1**: Thorax length of ACO and CO selection regime females measured to compare the body size of experimental females. Dot represents the mean across all four replicate populations and error bars represents standard error of mean, standard errors are calculated considering block means. Effect of selection regime was found to be significant on thorax length of selection regime females.

**S2: Results of mortality analysis for confounding effect of body size of females**

To analyse the confounding effect of body size on female mortality, the mortality analysis was performed in two ways a) without considering body size as covariate and b) considering body size as covariate. The mortality data was analysed as GLMM with poisson distribution. Here the block mean values of prop. dead under CE condition was converted into success and trials and the trials were included as offset in the model. Thorax length of females obtained by method described in supplementary information above was taken as covariate and selection regime (two levels: ACO and CO) as fixed factor. The block mean values were taken as unit of analysis. Block was taken as random factor.

Following model was used for the generalised linear mixed-effect model in R version 4.2.1 (R core Team, 2022) using glmmTMB package.

a) With body size as covariate

m1<-glmmTMB(Success ~ thorax_length+selection regime + (1|Block)+offset(log(Trial)), datafile, family = poisson(link = "log"))

| **Effect** | **Chisq** | **df** | **Pr(>Chisq)** |
| --- | --- | --- | --- |
| Thorax length | 8.2101 | 1 | **0.004** |
| Selection regime | 38.1189 | 1 | **<0.001** |

**Table S1**: Summary of results of generalized linear mixed-effect model (GLMM) on female mortality under CE assay condition. ANOVA table is obtained by Type III Wald chisquare tests. Selection regime was modelled as fixed factor and thorax length as covariate. All tests were done considering α=0.05 and significant p-values are mentioned in bold font style.

b) Without body size as covariate

m2 <- glmmTMB(Success ~selection regime+(1|Block)+offset(log(Trial)), datafile, family = poisson(link = "log"))

| **Effect** | **Chisq** | **df** | **Pr(>Chisq)** |
| --- | --- | --- | --- |
| Selection regime | 10668.835 | 1 | **<0.001** |

**Table S2**: Summary of results of generalized linear mixed-effect model (GLMM) on female mortality under CE assay condition. ANOVA table is obtained by Type III Wald chisquare tests. Selection regime was modelled as fixed factor and thorax length as covariate. All tests were done considering α=0.05 and significant p-values are mentioned in bold font style.

**S3: Results of block-wise analysis on age-specific per capita fecundity**

To analyse block-wise data on age-specific per capita fecundity, following model was used for the linear mixed-effect model in R version 4.2.1 (R Core Team, 2022) using lme4 package and lmerTest:

Per capita fecundity ~ Selection regime*Age+ (1|Block/Vial_ID)

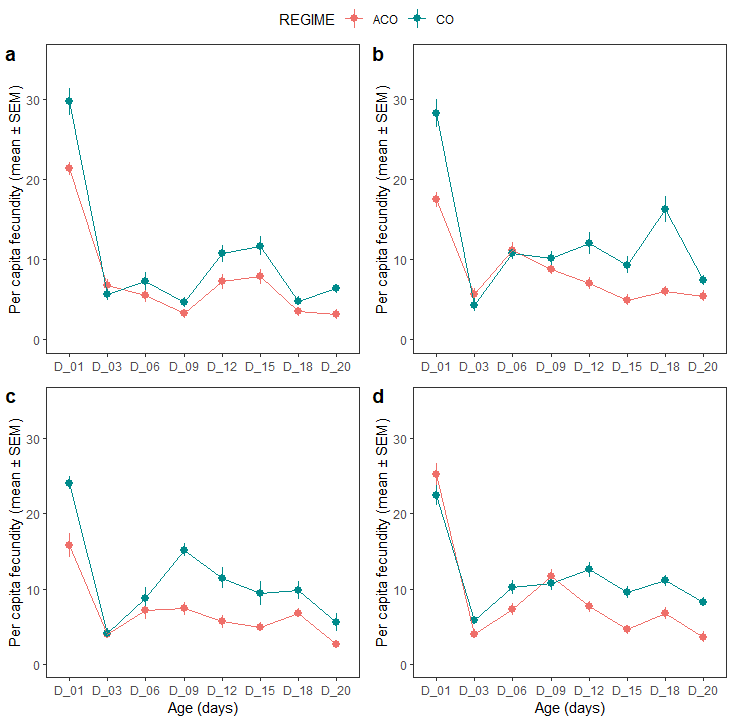

**Figure S3**: Age specific per capita fecundity across ACO and CO selection regime females held with ancestral CO males under continuous exposure (CE) assay condition for 20 days. Dot represents the mean and error bars are standard error. Effect of selection regime, age and selection regime × age was found to be significant on age specific per capita fecundity. The panels show results from (a) Block 1, (b) Block 2, (c) Block 3, and (d) Block 4.

| **Effect** | **Block** | **SS** | **MS** | **Num DF** | **Den DF** | **F** | **p** |
| --- | --- | --- | --- | --- | --- | --- | --- |
| Selection regime | **1** | 164.0 | 163.99 | 1 | 27.803 | 16.771 | **<0.001** |
| Age |  | 10572.1 | 1510.31 | 7 | 194.006 | 154.463 | **<0.001** |
| Selection regime × age |  | 393.7 | 56.25 | 7 | 194.006 | 5.753 | **<0.001** |
| Selection regime | **2** | 278.3 | 278.28 | 1 | 27.890 | 23.863 | **<0.001** |
| Age |  | 6384.1 | 912.01 | 7 | 194.970 | 78.206 | **<0.001** |
| Selection regime × age |  | 1057.6 | 151.08 | 7 | 194.970 | 12.955 | **<0.001** |
| Selection regime | **3** | 17.043 | 17.0431 | 1 | 26.000 | 37.229 | **<0.001** |
| Age |  | 124.137 | 17.7338 | 7 | 182.000 | 38.737 | **<0.001** |
| Seection regime × age |  | 8.182 | 1.1688 | 7 | 182.000 | 2.553 | **0.016** |
| Selection regime | **4** | 5.641 | 5.6409 | 1 | 28.000 | 21.644 | **<0.001** |
| Age |  | 148.212 | 21.1731 | 7 | 196.000 | 81.240 | **<0.001** |
| Selection regime×age |  | 13.001 | 1.8572 | 7 | 196.000 | 7.126 | **<0.001** |

**Table S2**: Summary of results of linear mixed-effect model (LMM) on female age-specific per capita fecundity under CE assay condition for each block separately. Selection regime, age and their two-way interaction were modelled as fixed factors. All tests were done considering α=0.05 and significant p-values are mentioned in bold font style.

**S4: Results of day1 per capita fecundity analysis**

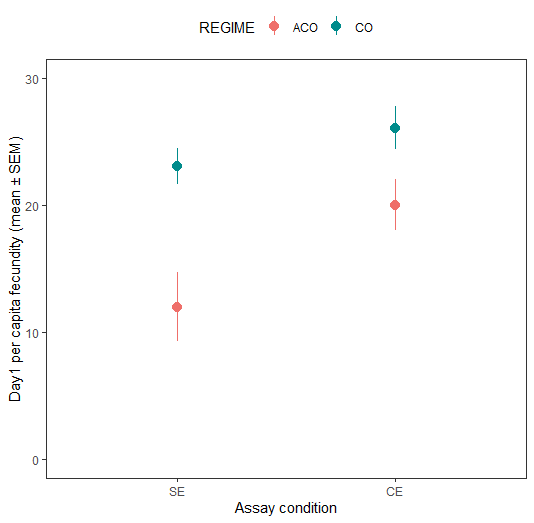

**Figure S4**: Day1 per capita fecundity across ACO and CO selection regime females held with ancestral CO males under single exposure (SE) and continuous exposure (CE) assay. Dot represents the mean across all four replicate populations and error bars represents standard error of means, standard errors are calculated considering block means. Effect of selection regime and assay condition was found to be significant on day1 age specific per capita fecundity.

**S5: Results of the test of random effect of block**

| **Trait** | **Effect** | **npar** | **AIC** | **LogLik** | **Chisq** | **DF** | **Pr(>Chisq)** |
| --- | --- | --- | --- | --- | --- | --- | --- |
| **Cumulative per capita fecundity** | Block × Selection regime × Assay condition | 8 | 1913.40 | -948.71 | 3.092 | 1 | 0.079 |
|  | Block × Assay condition | 8 | 1910.30 | -947.17 | 0.015 | 1 | 0.902 |
|  | Block × Selection regime | 8 | 1910.60 | -947.29 | 0.257 | 1 | 0.612 |
|  | Block | 8 | 1910.60 | -947.29 | 0.239 | 3 | 0.625 |
| **Day 1 per capita fecundity** | Block × Selection regime × Assay condition | 8 | 1501.50 | -742.74 | 1.167 | 1 | 0.280 |
|  | Block ×Assay condition | 8 | 1500.70 | -742.36 | 0.404 | 1 | 0.525 |
|  | Block × Selection regime | 8 | 1502.40 | -743.22 | 2.127 | 1 | 0.145 |
|  | Block | 8 | 1500.30 | -742.16 | 0 | 1 | 1 |
| **Age specific per capita fecundity** | Block × Selection regime × Age | 21 | 1904.50 | -931.24 | 20.803 | 1 | **<0.001** |
|  | Block × Age | 21 | 1892.50 | -925.24 | 8.817 | 1 | **0.003** |
|  | Block × Selection regime | 21 | 1883.70 | -920.83 | 0 | 1 | 1 |
|  | Block | 21 | 1884.00 | -920.97 | 0.278 | 1 | 0.598 |
| **Body size** | Block × Selection regime | 4 | -1029.30 | 518.64 | 0 | 1 | 1 |
|  | Block | 4 | -1026.70 | 517.36 | 2.542 | 1 | 0.111 |

**Table S3**: Effect of block as a random factor and those of all two-way interactions involving block on cumulative per capita fecundity, day 1 per capita fecundity, age specific per capita fecundity and body size. This was done using a linear mixed-effect model in R version 4.2.1 using lme4 package (R Core Team, 2022) and lmerTest. Significant effects (p < 0.05) are shown in bold font style

**S6: Models used for the main analysis**

Following model was used for the linear mixed-effect model in R version 4.2.1 using lme4 package (R Core Team, 2022) and lmerTest:

| **Trait** | **Model** |
| --- | --- |
| Cumulative female fecundity (CUF) | CUF ~ Selection regime × assay condition + (1\|Block) + (1\|Block:Selection regime) + (1\|Block:assay condition) + (1\|Block:selection regime:assay condition) |
| Age-specific per capita fecundity (PCF) | PCF ~ Selection regime × Age + (1\|Block/Vial_ID) + (1\|Block:Selection regime) + (1\|Block:Age) + (1\|Block:Selection regime:Age) |
| Day 1 per capita fecundity (PCFD1) | PCFD1 ~ Selection regime × assay condition + (1\|Block) + (1\|Block:Selection regime) + (1\|Block:assay condition) + (1\|Block:Selection regime:assay condition) |
| Thorax length (THXL) | THXL ~ Selection regime + (1\|BLOCK) |
